# Visual features are processed before navigational affordances in the human brain

**DOI:** 10.1101/2023.06.27.546695

**Authors:** Kshitij Dwivedi, Sari Sadiya, Marta P. Balode, Gemma Roig, Radoslaw M. Cichy

## Abstract

To navigate through their immediate environment humans process scene information rapidly. How does the cascade of neural processing elicited by scene viewing to facilitate navigational planning unfold over time? To investigate, we recorded human brain responses to visual scenes with electroencephalography (EEG) and related those to computational models that operationalize three aspects of scene processing (2D, 3D, and semantic information), as well as to a behavioral model capturing navigational affordances. We found a temporal processing hierarchy: navigational affordance is processed later than the other scene features (2D, 3D, and semantic) investigated. This reveals the temporal order with which the human brain computes complex scene information and suggests that the brain leverages these pieces of information to plan navigation.

## Introduction

By looking even only briefly at a scene, we rapidly extract multifaceted pieces of visual information^1–4^ that enable us to navigate through the scene, for example by planning a route through it. How does the brain compute visual information that affords navigational route planning in a scene?

Identifying navigational affordances is a complex computational feat that requires the processing of several different scene features, such as 3-dimensional and semantic scene aspects. For instance, successfully navigating the immediate environment requires localizing obstacles and finding out a way around them, which necessitates 3D scene information. Similarly, semantic scene classification can also benefit route planning as navigating typical basements, balconies, and garages require different procedures. We thus hypothesized that visual features crucial for identifying navigational affordances are processed before processing navigational affordances.

To test the hypothesis, we collected human electroencephalography (EEG) responses to indoor scene images, capturing the temporal order of scene feature processing in the human brain during visual scene perception.

We investigated three types of visual features: 2-dimensional (2D), 3-dimensional (3D) and semantic features. We operationalized the visual features in the indoor scene images as activations of deep neural networks (DNNs) trained to perform respective 2D, 3D and semantic tasks^5^. Navigational features are captured using navigational affordance maps (NAM) constructed using human behavioral responses when planning exit routes in natural indoor scene images^6^.

We then related the visual and navigational features to EEG data using representational similarity analysis (RSA) (Kriegeskorte et al., 2008) in a time-resolved manner, yielding time courses with which visual representations of particular features emerge. Finally, we compared these features and NAM with EEG, revealing the temporal order in which these features are processed in the human brain.

We found that navigational affordance representations emerged significantly later than visual features. This is consistent with the view that the brain uses 2D, 3D and semantic scene features to facilitate navigation planning.

## Results

We recorded EEG responses from 16 healthy volunteers (7 females, mean age 28.9 ± SD 5.6) to 50 indoor scene images. While viewing the stimuli, participants were asked to assess navigational affordance by imagining the directions of the navigational paths relative to the participant’s viewpoint, i.e., whether the paths were leading to the left, the center, or the right (Figure 1A).

**Figure 1.**
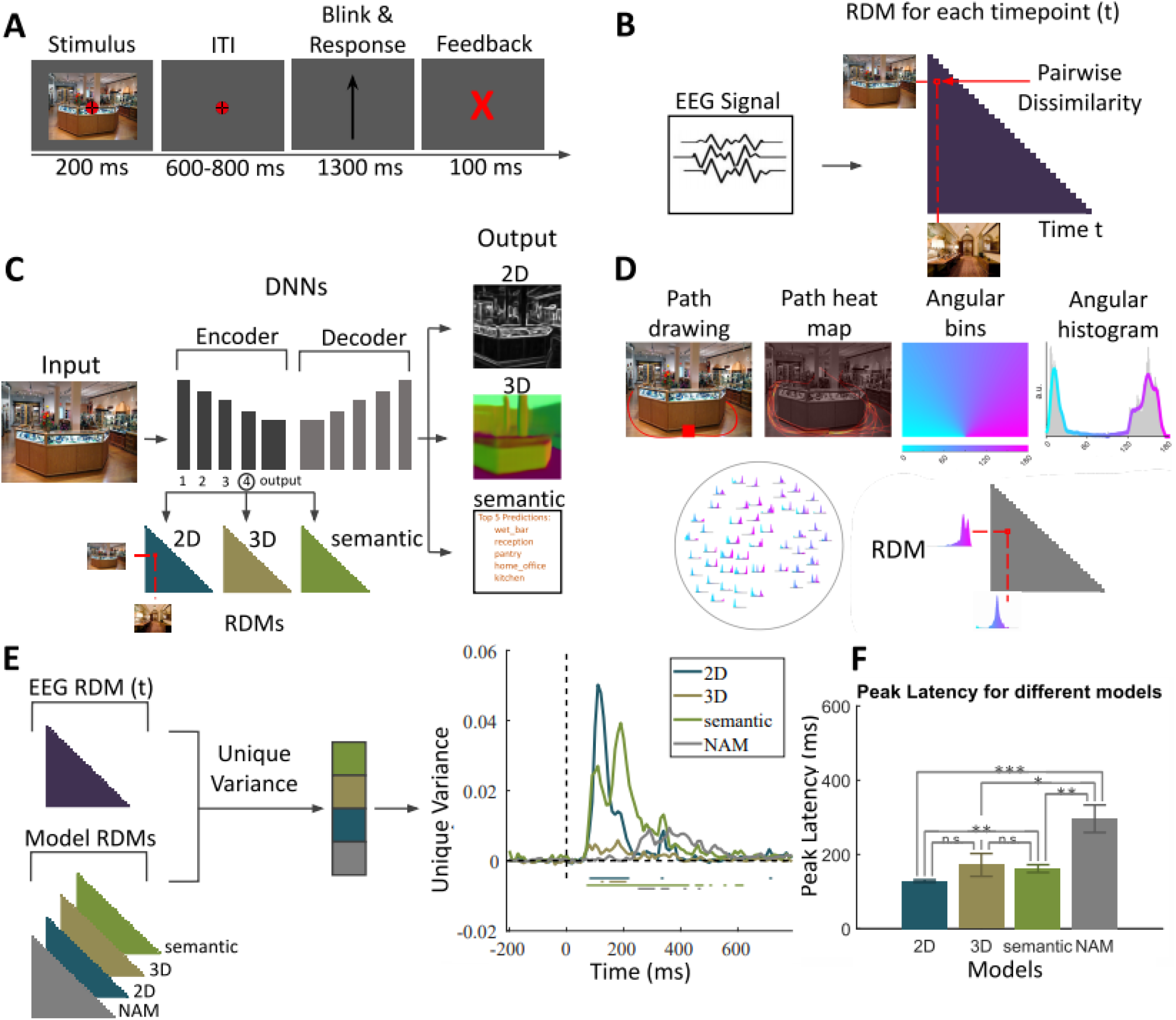
A) EEG paradigm. Participants viewed 50 images of indoor scenes and were asked to mentally plan possible exit paths through the scenes. On interspersed catch trials participants had to respond whether the exit path displayed on the screen corresponded to any of the exit paths from the previous trial. B) EEG RDMs. We computed RDMs for each EEG time point (every 10 ms from -200 to +800 ms with respect to image onset). DNN RDMs. We calculated RDMs from the activations extracted from the 4th block and output layer of a ResNet50 DNN trained on 2D, 3D and semantic tasks. D) NAM model and RDM (Bonner and Epstein, 2018). E) Variance partitioning. We calculated the unique EEG variance explained by each of the models, revealing different temporal activation patterns. Lines below the plots indicate significant times using t-test (FDR corrected p < 0.05). F) Peak latencies of different models. Bars indicate the peak latencies for different models. Error bars indicate standard deviation for 16 subjects. Stars above bars indicate significant differences across different models (*p <0.05,**p < 0.01, ***p < 0.001, t-test FDR corrected).

We then investigated when representations of visual and navigational features emerge in the human brain by comparing the EEG data to deep neural network (DNN) models and behavioral data operationalizing those features using representational similarity analysis.

For this we first transformed the peri-stimulus EEG responses (from -100 to +800ms with respect to stimulus onset) into representational dissimilarity matrices (RDMs) (Figure 1B) in steps of 10ms. We then created the 2D, 3D, and semantic RDMs using the activations of DNNs trained on 2D, 3D, and semantic tasks (Figure 1C).

To construct the navigational affordance model RDM, subjects were asked to indicate the exit routes starting from the bottom of a scene image presented to them using a computer mouse. Then, probabilistic heatmaps of navigational affordances were created pooling e data across subjects. These heatmaps were transformed into angular histograms that approximate a probabilistic navigational affordance map (NAM) of potential navigational paths radiating from the viewer’s perspective. Pairwise comparison between NAMs resulted in NAM RDMs (Figure 1D).

Having transformed all the modalities into a common representational space, we performed variance partitioning via regression to find out how much variance of an EEG RDM at a given time point is explained uniquely by the RDMs of any given model (Figure 1E, left panel). For this, we first performed a regression with all model RDMs as the independent variables and EEG RDM as the dependent variable. This determined the variance explained by all the models together 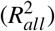. Then, we performed a second set of regressions, removing the RDMs of a given model (e.g., NAM) from the independent variables to find the variance explained by models leaving out the model of interest 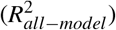. The unique variance of the EEG RDM explained by the selected model is then calculated as 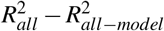.

We found all models explained unique variance in EEG, but to different degrees (Figure 1E, right panel). The 2D DNN RDM explained the most variance in EEG (max R2=0.0502), followed by the semantic DNN (max R2=0.0393). The contributions of NAM and 3D DNN RDMs were lower (NAM RDM max R2=0.0094, 3D DNN RDM max R2=0.0058). This provides evidence that all feature representations can be uniquely tracked in our experiment, and allowed us to inspect the time course further.

We observed a temporal pattern in peak timings (Figure 1F). The highest uniquely explained variance by the 2D DNN RDM occurred first at 128.12±3.56ms after stimulus onset, followed by 3D and semantic DNN RDMs peaking at 171.87±30.79ms and 161.87±10.45ms, respectively. The unique variance of the NAM RDM reached its peak significantly later than the other model RDMs at 296.25±37.05ms after stimulus onset. This suggests a hierarchy of scene feature processing leading up to the representation of navigational affordances.

## Discussion

In this study, we investigated the temporal dynamics of scene perception, focusing particularly on the temporal order in which 2D, 3D, semantic features and navigational affordances emerge. We found that the emergence of 2D, semantic and 3D features preceded the emergence of navigational affordance representations.

The early emergence of low-level 2D features followed by high-level semantic features has been previously observed^7^, in particular in studies investigating the correspondence of layers of DNNs trained on scene classification tasks with time-resolved human EEG responses^8^. Equating early layers with low-level visual features and later layers with high-level features yielded a temporal hierarchy, as also observed for object processing^9^. In contrast, 3D features have attracted less attention in M/EEG studies and thus the temporal dynamics with which 3D feature representations emerge are less well understood. Although some studies investigated the temporal dynamics of spatial layout^10,11^, spatial layout represents only coarse-grained 3D features such as the size of the scene in the real world and the position of large surfaces, but does not take into account fine-grained 3D information such as the pose of different objects present in the scene. fMRI studies have investigated fine-grained spatial 3D features by investigating the representation of surface normals^12^ and correspondence to DNNs trained to solve 3D tasks (e.g. depth, occlusion) on the Taskonomy dataset^13,14^. Here we complement these efforts in the temporal dimension by showing that 3D features are processed in parallel with semantic features.

Navigational affordance representation emerged significantly later than 2D, 3D, and semantic representations. This suggests by temporal order that humans leverage those features to process navigational affordances. In contrast to our results, a recent study^15^ reported both early and late physiological markers of navigational affordances. We note the key differences in both the studies that may have led to different conclusions about navigational affordance processing. First, the images used here are natural and complex, while in^15^ the images were simple and synthetic. Image complexity can influence processing time, potentially due to recurrence^16^. Second, in our study, subjects were tasked to identify and find their way around obstacles, making them process occlusions and 3D scene information. In Harel et al. (2022), subjects had to count the number of doors, for which processing 3D scene information might not be needed^15^. The difference in timing might thus reflect a difference in feature processing as required by the task.

A limitation on the ecological validity of our study is that we used static images to assess temporal aspects of scene perception, whereas in real-world situations, humans would process moving images, especially while navigating through indoor environments. Another aspect that could be addressed in the future is inclusion of additional computational models in studying the temporal dynamics of scene perception. Nevertheless, our findings demonstrate a timeline of hierarchical scene feature processing, suggesting that the visual scene features investigated here support navigational planning.

## Methods

### EEG

#### Participants

We recorded EEG data from 16 healthy volunteers (7 females, mean age 28.9 ± SD 5.6). All participants had normal or corrected-to-normal vision. Participants gave informed consent before the experiment and were provided with monetary compensation. The experiment was conducted in compliance with the Declaration of Helsinki and was approved by the ethics committee of the Freie Universität Berlin.

#### Stimuli

The stimuli were 50 color images of different indoor environments with easily detectable navigational paths originating at the bottom center of each image, previously used in a study by Bonner and Epstein (2017)^17^. The dimensions of all the images were 1024 × 768 pixels and subtended 7° of visual angle in width and 5.25° in height. They were presented on a gray screen with a combination of bull’s-eye and crosshair fixation targets^18^ positioned centrally.

#### Experimental Paradigm

The paradigm was designed to engage the participants in explicit navigational affordance processing of every image. While viewing the stimuli, participants were asked to imagine the directions of the navigational paths relative to the participant’s viewpoint, i.e., whether the paths were leading to the left, the center, or the right (Figure 1A).

On each trial Images were presented for 200ms followed by a randomly varying inter-trial interval between 600ms and 800ms. Image presentation trials were ordered in blocks of one to 5 trials in length.

Blocks were followed by the presentation of a catch trial during which participants had to conduct a task meant to ensure that participants remained attentive and processed the images with respect to spatial and navigational aspects. During catch trials an arrow appeared on the screen for 1.3s, during which the participants had to indicate whether an arrow on the screen pointed in the same (congruent) or in a different (incongruent) direction than the navigational path in the previous trial. Participants had to respond by pressing the right arrow key for “yes” (congruent, pointing in the same direction) and the left arrow key for “no” (incongruent, not pointing in the same direction). After the response, feedback was presented for 0.1s, followed by a post-feedback time of 0.2s. The number of congruent and incongruent trials was balanced across the experiment.

Blocks were organized in runs: there were 69 blocks in total (24 blocks of 5 trials, 15 blocks of 3 and 15 blocks of 4 trials, 10 blocks of 2 trials, and 5 of 1 trial) for each run presented in random order. There were 15 runs (6.2 minutes each) in total in the experiment. This design resulted in each image being repeated 75 times.

#### EEG Recording and pre-processing

For all participants, we recorded continuous neural activity with EEG using the Easycap 64-channel standard electrode system and Brainvision actiCHamp amplifier. We followed the 10-10 system for electrode placement. EEG signals were recorded with a sampling rate of 1000Hz and bandpass filtered online between 0.03Hz and 100Hz.

All electrodes were online referenced to the FCz electrode and grounded to the AFz electrodes. We pre-processed the data offline using FieldTrip^19^ and selected 17 posterior and occipital channels for further analysis. The data was segmented in epochs of -0.2s to 0.8s relative to stimulus onset, baseline-corrected to the average pre-stimulus signal, and down-sampled to 100Hz. Eye blinks and other artifacts were identified with independent component analysis and removed after inspection.

#### Pairwise Decoding

To determine how well EEG signals can be used to differentiate between the 50 scene images, we calculated the pairwise decoding accuracy score for each image pair at every time point using CoSMoMVPA^20^. This was done in a time-resolved manner, assessing 100 time points every 0.01s from -0.2 to 0.8s relative to image onset. For every possible pair of image conditions, we partitioned the pre-processed EEG signals across all trial repetitions into training and test data using a leave-one-trial-out cross-validation scheme. We then trained LDA classifiers on all-but-one trials and tested them on the left-out trials. Decoding accuracy scores were averaged across cross-validation folds. To create a grand average time series of EEG decoding accuracy, we calculated the mean decoding accuracy across all pairs.

### Behavioral Data

To quantify the navigational affordance features in the 50 experimental images, we used the navigational affordance model (NAM) by Bonner and Epstein (2017)^17^, which was created using the same set of 50 images. Bonner and Epstein asked participants to draw all possible navigational paths in each image starting from the bottom center of the image. The responses of the participants were aggregated together into heatmaps. Then angular binning was performed to create a navigational affordance histogram by counting the number of pixels in each one-degree bin from 0-180°.

### Deep Neural Network models

To assess low-, mid-, and high-level features of indoor scenes, we used activations from 18 pre-trained deep neural network (DNN) models from the Taskonomy Task Bank^5^. The Task Bank consists of DNN models that cover various tasks in computer vision, which can be grouped into three categories (2D, 3D, and semantic tasks). Models from the 2D task category process low-level visual features (for example, 2D edges and key points), models from the 3D category process such mid-level features as surface normals and depth, whereas models trained on classification tasks process high-level semantic features.

All models were trained on the Taskonomy dataset^5^, which consists of 4.5 million fully annotated images of indoor environments from 600 buildings. The model architecture is built on an encoder and a decoder. The encoder is identical across the 18 task models, being based on ResNet-50^21^ with a compressed convolutional output layer. The decoder architecture differs between the models; for example, 2D and 3D models have a decoder structure of 15 convolution and deconvolution layers, whereas the scene classification model has only two fully connected layers in the decoder because of its low output dimensions.

To ensure comparability across models, for each model, we selected the block4 and the output layer from the encoder as the representative task-specific layers following^13^.

### EEG-DNN/Model comparison

To compare the EEG responses with the DNN and behavioral responses we used representational similarity analysis^22^. In RSA, data from different incommensurate sources are related using a common summary of the representational geometry of each source. For this we first computed representational dissimilarity matrices (RDMs) for each model and for the EEG data. RDMs are diagonally symmetric square NxN dimensional matrices (where N is the number of conditions) that summarize the dissimilarity between condition-specific responses in each source space.

Then, we performed a variance partitioning analysis^23^ to estimate the unique variances of the EEG RDMs explained by the model RDM investigated in this work. We detail the RDM construction below.

#### EEG RDMs

We used pairwise decoding accuracy between image conditions to construct EEG RDMs. The rationale is that the more dissimilar the EEG signals arising from two different images are, the higher the decoding accuracy score for that pair of images will be. RDMs were constructed in a time-resolved manner for each time point (Figure 1 B), yielding 50×50 EEG RDM for each of the 100 time points.

#### Model RDMs

The NAM RDM was constructed by computing the euclidean distance between navigational affordance histograms of all pairs of images (Figure 1D). We downloaded the precomputed NAM RDMs from https://figshare.com/s/5ff0a04c2872e1e1f416.

For each DNN RDM, we selected block4 and the encoder output layer for creating RDMs. We measured the dissimilarity between any two image representations in the DNN by calculating 1 minus the Pearson correlation distance (1-*ρ*) between the corresponding layer activations. This resulted in two 50×50 RDMs for each DNN (i.e., the block4 and the output layer RDM). RDMs were aggregated and averaged by DNN group for each task type (2D, 3D, semantic) (Figure 1C), resulting in two RDMs for each task type.

#### Variance Partitioning analysis

Since the DNNs investigated are trained on the same dataset, their RDMs are expected to be correlated. To nevertheless identify the aspects unique to a particular task type, we use variance partitioning with the goal to identify variance uniquely attributable to any one model. To compute the unique variance of a given model, we calculated the difference in variance explained when all the model RDMs are used as independent variables and variance explained when all but current model RDMs are used as independent variables.

We conducted this analysis at every time point separately, i.e. every 10ms from -200 to +800ms relative to image onset. In the regression we used the lower triangular part of the RDM as it describes the representational geometry fully and avoids potential artifacts created by including the diagonal. This resulted in four time series, one for each model (3D, 3D, semantic, navigational affordance) indicating when feature representations corresponding to the model type emerge during visual processing.

#### Statistical analysis

We applied a two-sided t-test to assess the statistical significance of the unique variance explained by different models and peak latencies. We corrected the p-values for multiple comparisons by applying FDR correction with a threshold of 0.05.

## Data and Code Availability

The code and data necessary for reproducing the results presented in this paper can be found at https://osf.io/wz4ha/.

## Acknowledgments

This work was supported by the German Research Council grants (CI241/1-1, CI241/3-1, CI241/7-1) awarded to Radoslaw Cichy, the German Research Council grants (FOR 5368 ARENA, RO 6458/2-1, RO 6458/2-1) awarded to Gemma Roig, and a European Research Council grant (ERC-StG-2018-803370) awarded to Radoslaw Cichy. Additional support was provided by the Hessian ministry of Art and Science LOEWE program through the Frankfurt Center for Multi-scale Modeling in the Life Sciences. Computing resources were provided by the high-performance computing facilities at ZEDAT, Freie Universität Berlin^24^.

## Author contributions statement

Must include all authors, identified by initials, for example: A.A. conceived the experiment(s), A.A. and B.A. conducted the experiment(s), C.A. and D.A. analysed the results. All authors reviewed the manuscript. K.D. worked on this manuscript prior to joining Amazon Inc.

## Notes

https://osf.io/wz4ha/

### Competing Interest Statement

The authors have declared no competing interest.

### Summary of Updates

Formatting, using late template instead of word

## References

1. Fei-Fei, L., Iyer, A., Koch, C. & Perona, P. What do we perceive in a glance of a real-world scene? J. Vis. 7(1), 10, DOI: https://doi.org/10.1167/7.1.10 (2007).

2. Greene, M. R. & Oliva, A. The briefest of glances: The time course of natural scene understanding. Psychol. Sci. 20(4), 464–472 (2009).

3. Potter, M. C. Meaning in visual search. Science 187(4180), 965–966, DOI: https://doi.org/10.1126/science.1145183 (1975).

4. Thorpe, S., Fize, D. & Marlot, C. Speed of processing in the human visual system. Nature 381(6582), 520–522., DOI: https://doi.org/10.1038/381520a0 (1996).

5. Zamir, A. R. et al. Taskonomy: Disentangling task transfer learning. Proc. IEEE conference on computer vision pattern recognition 3712–3722. (2018).

6. Bonner, M. F. & Epstein, R. A. Computational mechanisms underlying cortical responses to the affordance properties of visual scenes. PLoS Comput. Biol. 14(4), e1006111 (2018).

7. Harel, A., Groen, I. I. A., Kravitz, D. J., Deouell, L. Y. & Baker, C. I. The temporal dynamics of scene processing: A multi-faceted eeg investigation. ENeuro DOI: https://doi.org/10.1523/ENEURO.0139-16.2016 (2016).

8. Greene, M. R. & Hansen, B. C. Shared spatiotemporal category representations in biological and artificial deep neural networks. PLOS Comput. Biol. 14(7), e1006327, DOI: https://doi.org/10.1371/journal.pcbi.1006327 (2018).

9. Cichy, R. M., Khosla, A., Pantazis, D., Torralba, A. & Oliva, A. Comparison of deep neural networks to spatio-temporal cortical dynamics of human visual object recognition reveals hierarchical correspondence. Sci. Reports 6, 27755 (2016).

10. Cichy, R. M., Khosla, A., Pantazis, D. & Oliva, A. Dynamics of scene representations in the human brain revealed by magnetoencephalography and deep neural networks. NeuroImage 153, 346–358 (2017).

11. Henriksson, L., Mur, M. & Kriegeskorte, N. Rapid invariant encoding of scene layout in human opa. Neuron (2019).

12. Lescroart, M. D. & Gallant, J. L. Human scene-selective areas represent 3d configurations of surfaces. Neuron 101(1), 178–192 (2019).

13. Dwivedi, K., Bonner, M. F., Cichy, R. M. & Roig, G. Unveiling functions of the visual cortex using task-specific deep neural networks. PLOS Comput. Biol. 17(8), e1009267., DOI: https://doi.org/10.1371/journal.pcbi.1009267 (2021).

14. Wang, A. Y., Wehbe, L. & Tarr, M. J. Neural taskonomy: Inferring the similarity of task-derived representations from brain activity. Adv. Neural Inf. Process. Syst. 32 DOI: https://doi.org/10.1101/708016 (2019).

15. Harel, A., Nador, J. D., Bonner, M. F. & Epstein, R. A. Early electrophysiological markers of navigational affordances in scenes. J. Cogn. Neurosci. 34(3), 397–410, DOI: https://doi.org/10.1162/jocn_a_01810 (2022).

16. Kar, K., Kubilius, J., Schmidt, K., Issa, E. B. & DiCarlo, J. J. Evidence that recurrent circuits are critical to the ventral stream’s execution of core object recognition behavior. Nat. Neurosci. 22(6), 974–983, DOI: https://doi.org/10.1038/s41593-019-0392-5 (2019).

17. Bonner, M. F. & Epstein, R. A. Coding of navigational affordances in the human visual system. Proc. Natl. Acad. Sci. 114(18), 4793–4798 (2017).

18. Thaler, L., Schütz, A. C., Goodale, M. A. & Gegenfurtner, K. R. What is the best fixation target? The effect target shape on stability fixational eye movements. Vis. Res. 76, 31–42, DOI: https://doi.org/10.1016/j.visres.2012.10.012 (2013).

19. Oostenveld, R., Fries, P., Maris, E. & Schoffelen, J.-M. Fieldtrip: Open source software for advanced analysis of meg, eeg, and invasive electrophysiological data. Comput. Intell. Neurosci. 2011, e156869, DOI: https://doi.org/10.1155/2011/156869 (2010).

20. Oosterhof, N. N., Connolly, A. C. & Haxby, J. V. Cosmomvpa: Multi-modal multivariate pattern analysis of neuroimaging data in matlab/gnu octave. Front. Neuroinformatics 10 (2016).

21. He, K., Zhang, X., Ren, S. & Sun, J. Deep residual learning for image recognition. In 2016 IEEE Conference on Computer Vision and Pattern Recognition (CVPR), 770–778, DOI: 10.1109/CVPR.2016.90 (2016).

22. Kriegeskorte, N., Mur, M. & Bandettini, P. A. Representational similarity analysis-connecting the branches of systems neuroscience. Front. Syst. Neurosci. 2, 4 (2008).

23. Legendre, P. Studying beta diversity: Ecological variation partitioning by multiple regression and canonical analysis. J. Plant Ecol. 1(1), 3–8, DOI: https://doi.org/10.1093/jpe/rtm001 (2008).

24. Bennett, L., Melchers, B. & Proppe, B. Curta: A general-purpose high-performance computer at zedat, freie universität berlin, DOI: http://dx.doi.org/10.17169/refubium-26754 (2020).

